# Immunojanus Particles for low-volume and isolation-free unlabeled characterization of small Extracellular Vesicle in biofluids: Characterization of disease type by surface marker profiling

**DOI:** 10.1101/2024.08.17.607528

**Authors:** Sonu Kumar, John Alex Sinclair, Tiger Shi, Han-Sheng Chuang, Satyajyoti Senapati, Hsueh-Chia Chang

## Abstract

Small extracellular vesicles (sEVs) are vital for cellular communication and serve as critical biomarker carriers for diseases such as cancer. However, quantifying and profiling sEV surface markers presents significant challenges due to the low concentration of specific sEV-bound proteins and interference by more abundant dispersed proteins. This paper presents Immunojanus Particles (IJPs), a new method that enables the direct detection of sEVs in less than an hour without isolation. The design of IJPs incorporates fluorescent and non-fluorescent halves, utilizing rotational Brownian motion to detect captured sEVs through the change in the blinking rate, without interference from the smaller dispersed proteins. We demonstrate a detection limit of 2E5 sEVs/mL with low sample volumes and the capability to characterize sEVs directly from plasma, serum, cell culture media, and urine. In a small pilot study involving 87 subjects, including individuals with colorectal cancer, pancreatic ductal adenocarcinoma, glioblastoma, Alzheimer’s disease, and healthy controls, our method accurately identified the type of disease with a high 0.90-0.99 AUC in a blind setting. Compared with an orthogonal ultracentrifugation plus surface plasmon resonance (UC+SPR) method that requires about 24 hours, the sensitivity and dynamic range of IJP are better by 2 logs.

## Introduction

Small Extracellular Vesicles (sEVs) are lipid-bilayer enclosed particles containing important cargo, including protein and nucleic acids, depending on their nature of biogenesis and cellular origin^1–6^. Secreted by cells into the extracellular matrix, they often contribute to cell-to-cell paracrine communication over a short distance^7–12^. However, many find their way into human plasma, potentially offering avenues for surveying the cellular landscape with a simple blood draw^13–17^. The extracellular matrix can especially become leaky for cancer and several other diseases to escalate sEV escape and disease dissemination far from the primary site^18,19^. This amplified release of sEVs into the microcirculation from diseased cells further enhances its prognosis and diagnosis potential^20–23^. Additionally, the unique surface markers on sEVs, which facilitate their uptake by specific cells—including the exchange between cancerous and healthy cells, present a novel strategy for targeted therapy^24–28^. By leveraging these markers, therapies can be designed to selectively deliver drugs directly to cancer cells, minimizing harm to healthy tissue. Therefore, profiling surface markers on small Extracellular Vesicles presents valuable opportunities across diagnostics, therapeutics, and drug delivery, underscoring its extensive potential in advancing medical science.

Quantifying surface markers on sEVs suffers from several challenges and limitations – these sEV surface markers can also appear in non-vesicular forms, such as soluble entities^29,30^. Often, only the vesicular proteins are enzymatically active or have higher activity and hence are the relevant markers^31–36^. Furthermore, these surface markers can be present alongside non-sEV proteins like albumin, which exists in concentrations over a billion-fold higher than sEVs in biofluids^29^ such as human plasma – non-specific adsorption of these non-sEV proteins can interfere with any immunoassay^37,38^. Thus, traditional methods frequently necessitate an sEV isolation step to eliminate surface markers in non-vesicular forms and non-sEV-associated contaminating proteins found in unprocessed biofluids^29,30,39^. Ultracentrifugation is the most common isolation step before any sample characterization. With its high capital cost and bulky instruments, this cumbersome and tedious process significantly hampers parallelization and field application. Moreover, the yield of sEV isolation cannot be consistent every time, thus introducing a yield bias due to isolation alone.

Even after isolating sEVs, other challenges complicate the characterization methods to permit the sub-picomolar detection limit necessary for sEV quantification. For example, after isolation, sEVs often aggregate^40^, thus requiring immediate characterization, while the isolation step creates lipoprotein aggregates that are difficult to distinguish from sEVs^41^. Given that sEVs are sized between proteins and cells, many techniques originally designed for proteins and cells—such as western blot and flow cytometry, respectively—have been adapted for sEVs^41–46^ without fully addressing their unique challenges. Techniques like western blot heavily depend on contamination-free samples, which are challenging to obtain even after extensive isolation steps^47–50^. Larger sEVs with higher cargo capacities can mask the cargo of the diagnostically relevant smaller sEVs, making protein-based sEV assays suffer from bias and reduced sensitivity. Meanwhile, flow cytometry struggles to detect particles below cellular dimensions; even state-of-the-art nano-flow cytometry faces difficulties identifying particles smaller than 100nm due to their low time of flight^51^, often requiring extensive labeling and a low limit of detection^51–54^. Other recently proposed methods include interferometry-based Exoview, which has only one log dynamic range, preventing it from studying less abundant sEV fractions and requiring extensive pretreatment and labeling ^55,56^. Other proposed methods in literature have reasonable sensitivity but require extensive labeling or pretreatment^57,58^. A gold-standard for sEV characterization, with standardized isolation and pretreatment procedures, is hence still unavailable.

Therefore, an isolation-free, sensitive, and rapid characterization method for sEVs is needed for biofluids, such as plasma, serum, urine, and even cell culture media, to be practical for clinical applications. Herein, we present the Immunojanus Particles (IJPs) that can profile sEVs without isolation and are less prone to interference. These IJPs are micron-sized spherical particles with one fluorescent half and one non-fluorescent half. The fundamental basis of detection relies on the rotational Brownian motion these particles undergo; a blinking effect similar to stars blinking in the night sky results from the thermal kicks that alternatively expose the fluorescent or non-fluorescent sides. The frequency of this blinking is highly sensitive, and the binding of any biological particle >50 nm can significantly affect its blinking rate, which is ideal for sEV detection. Abundant proteins like albumin and soluble versions of the targeted marker on sEVs are too small to produce any signal, allowing us to selectively quantify sEVs from plasma, serum, urine, and cell media using less than 10 microliters of sample volume with a limit of detection of ∼200 sEVs/uL. Additionally, because these beads are suspended in a solution, the long incubation period associated with mass transfer limited surface assays is shortened significantly as sEVs only need to diffuse to the nearest bead. Mass transfer limitation (faster kinetics than diffusion rate) ensures that the signal follows a universal scaling with all antibodies that satisfy the mass transfer limitation. This eliminates the variation that comes with the different affinity of antibodies from various sources, but also different affinities of the same antibody with a heterogeneous population of antibodies due to steric hindrance or avidity in a small subfraction of the sEVs, which means capture fraction is not representative of the bulk. Utilizing this unique feature, the IJP platform can circumvent issues often accompanied by varying antibody affinities and sources.

Using our IJP platform, we can screen and identify different diseases (in a blind setting) using sEV surface markers in a mixed pool of 87 human subjects with colorectal cancer, pancreatic ductal adenocarcinoma, glioblastoma, Alzheimer’s disease, and healthy subjects. We achieved high sensitivity and specificity without needing sample pretreatment and directly from human plasma in under 60 minutes. We have thoroughly benchmarked our platform against a day-long Ultracentrifugation (UC) and Surface Plasmon Resonance (SPR), which produced results consistent with IJPs but with reduced sensitivity and selectivity.

## Results and Discussion

### The Immunojanus Particle (IJP) Platform

We synthesized Immunojanus particles (IJPs) by creating a monolayer of fluorescent polystyrene particles on a glass slide. Following this, a gold layer approximately 10 nm thick was deposited onto the upper surface, giving the particles a Janus configuration with two distinct faces. Figure 1a provides a schematic depiction of the IJPs, illustrating micron-sized spheres with one hemisphere being fluorescent and the other coated with gold. This configuration produces a blinking fluorescence effect when observed under a microscope, as the non-fluorescent gold and fluorescent polystyrene alternate due to Brownian rotation.

**Figure 1:**
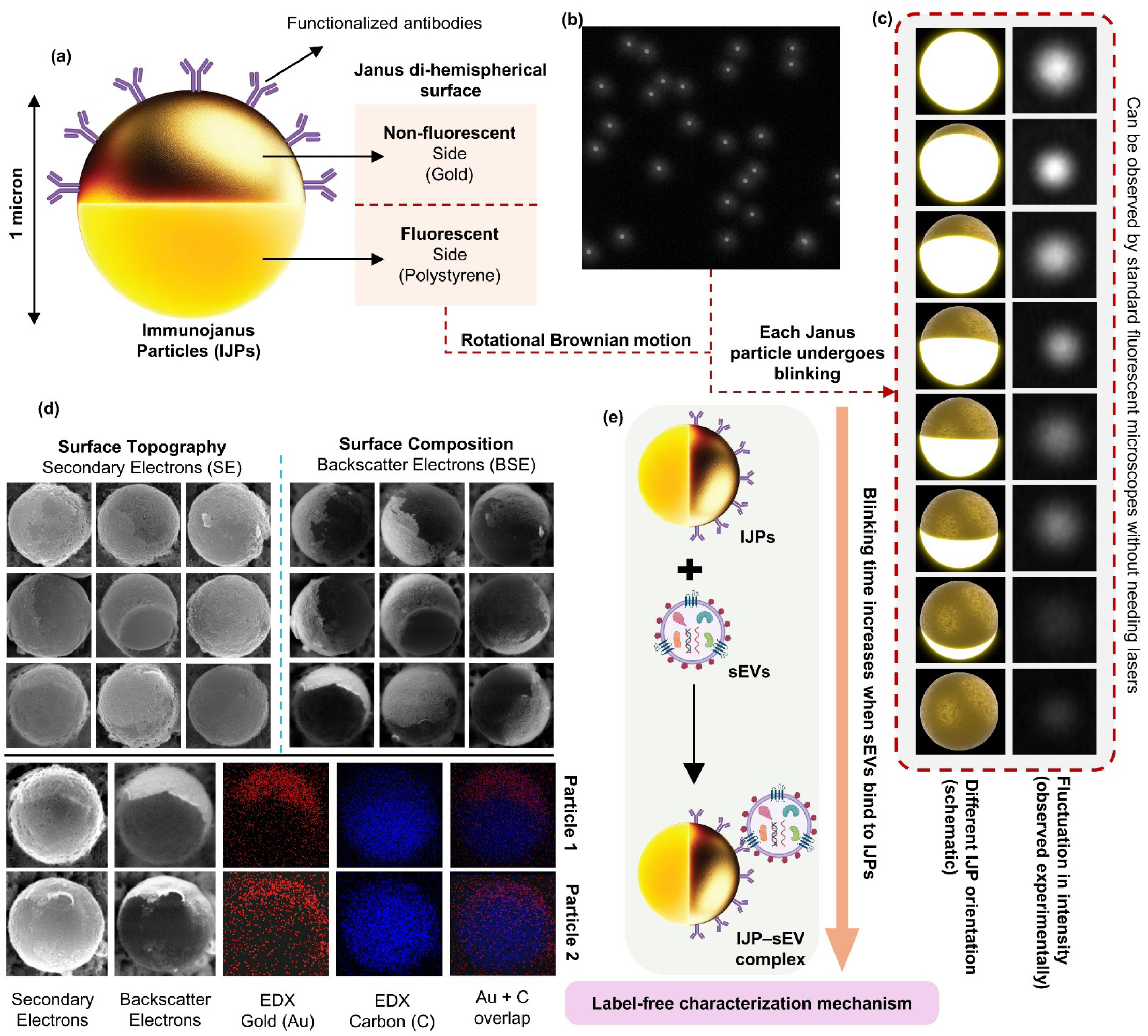
Immunojanus Particles are di-hemispherical micron-sized particles with one fluorescent side that makes them blink. The schematic (a) shows IJPs with an immunogold surface on one half (non-fluorescent) and a fluorescent half on the other, allowing the particle to switch between fluorescent and non-fluorescent states depending on orientation. A snapshot (b) captures dozens of IJPs in solution, each undergoing individual blinking. Due to rotational Brownian motion, different sides of the particles are oriented at different times, altering the intensity and causing the blinking effect (c). The left panel of (c) illustrates how different orientations result in high or low intensity, observed experimentally in the right panel. SEM images (d) reveal the spherical surface topography of IJPs using secondary electrons. Surface composition analysis using backscatter electrons shows one half of the particles covered with gold, consistent with the EDX detector findings, indicating gold (or a similar atomic number element) deposition on one half, making them appear brighter with backscatter electrons. (e) Schematic representation of the IJP sEV complex and its relative increase in time period compared to a blank IJP particle.

The gold hemisphere is functionalized with target-specific antibodies to capture relevant biological entities within a sample. Fig. 1b shows multiple Janus particles under a fluorescence microscope, and Fig. 1c illustrates their blinking behavior, typically about one second. SEM analysis in Fig. 1d reveals the topography and composition of the two hemispheres, highlighting their contrasting characteristics. EDX imaging confirmed the presence of a gold-like element on one hemisphere and a carbonaceous element similar to polystyrene on the other, demonstrating the Janus nature.

After functionalizing the gold hemisphere with antibodies, we allowed the IJPs to bind to antigens on sEVs that are specific to these antibodies. However, other nanocarriers, such as lipoproteins or soluble versions of these proteins can also bind to the antibodies on the IJP surface. Additionally, many non-specific proteins may bind non-specifically to the surface due to Van Der Waals forces, ionic interactions, and hydrophobic forces. This poses a significant challenge because our objective is to characterize sEVs without any isolation steps, such as Ultracentrifugation (UC), which typically removes these soluble and non-specific proteins. In the following sections, we will discuss the specificity of the signal and how our platform addresses these challenges. An important observation is that binding sufficiently large biological entities like sEVs to the IJPs significantly reduces the blinking frequency. This change is attributed to the increased drag exerted by the biological particle as it rotates within the surrounding fluid. Fig. 1e illustrates the mechanism employed for the IJPs signal, involving simple mixing and incubation procedures.

To accurately characterize sEVs, it is essential to determine the blinking periods of hundreds of IJPs per frame over extended durations. This requires the automated detection and tracking of IJPs within a frame, followed by calculating the average blinking frequency. For detection and tracking purposes, our focus is primarily on the change in intensity of the IJP fluorescence rather than high-definition imaging that reveals particle topology. High- resolution imaging typically depends on image segmentation and mask-based methods that utilize discontinuity detection or grayscale similarity and require expensive instrumentation for capturing high-resolution images. Moreover, incorporating more Janus particles per frame at lower magnification, instead of capturing high-resolution images of fewer IJPs at higher magnification, enhances our statistical analysis. This approach provides a more robust distribution of blinking periods by looking at the rate of change of particle intensity, leveraging the increased IJP sample size to improve the accuracy and reliability of our measurements while also requiring cheaper fluorescent optical systems. We report the change in blinking as the change in ensemble average of blinking rate across all IJPs.

We employ a distinct methodology for detecting IJPs and processing low-resolution images/videos, which involves multiplying the image by a dilation matrix corresponding to the feature size of JPs in these images to enlarge any high intensity point with a similar feature size. Post-dilation, we utilize a disk detection method based on accumulation points referred to as the Circular Hough Transform^59–61^ to detect the IJPs in the image, as shown in Fig. 2a. The detected centers are then mapped back onto the original image and tracked across subsequent frames. Key metrics, such as the intensity of the particle, are analyzed using wavelet techniques to determine the frequency of blinking. Larger aggregates, which do not blink due to their non-Janus nature and large size, are excluded by thresholding and focusing on the range of frequencies typical for Janus Particles. This algorithm and its applications are detailed in Fig. 2a, and all the intensities are measured by looking within the proximity of the detected center of the IJPs in the raw unprocessed footage.

**Figure 2:**
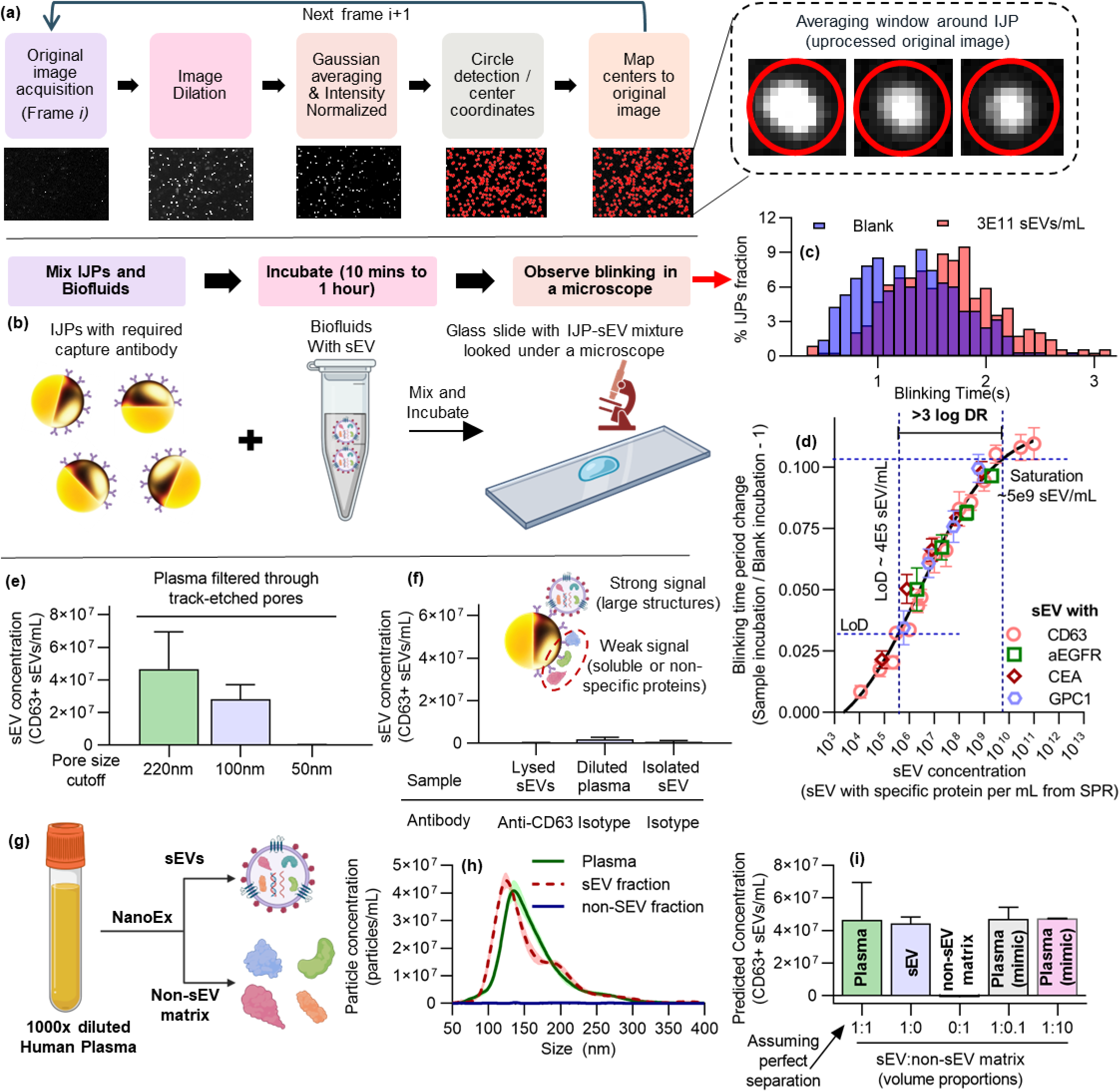
Characterization of small Extracellular Vesicles (sEVs) using Immunojanus Particles (IJPs). (a) Shows an automated algorithm for tracking individual IJPs, which involves image acquisition, dilation, Gaussian averaging, intensity normalization, circle detection, and mapping centers to the original image. (b) Illustrates the experimental process where IJPs with required capture antibodies are mixed with biofluids containing sEVs, incubated for 10 minutes to 1 hour, and observed under a microscope for blinking. (c) Histogram plot demonstrates the shift in the blinking time period of particles upon mixing with sEVs (3E11/mL) compared to blank (PBS). (d) Scatter plot provides a calibration curve for IJPs coated with anti-CD63, anti-aEGFR (mab806), anti-CEA, and anti-GPC1 against sEV concentration derived from surface plasmon resonance for isolated sEVs from DiFi cell line (n=2) and human plasma samples (n=3). Error bars represent standard error. (e) Bar plot shows signal variation from plasma filtered through 220nm, 100nm, and 50nm filters, indicating IJPs’ immunity to signals from species below 50nm. Most sEVs are 50-200nm, thus 100nm filtering reducing the signal, while 50nm suppresses it. Error bars represent one standard deviation. (f) Bar plot highlights that lysed sEVs (detergent-treated) with anti-CD63 capture, diluted plasma with isotype capture, and isolated sEV with isotype capture produce no signal. (g) Displays the process of splitting plasma into sEV and non-sEV matrix fractions to create mimic plasma by mixing varying proportions of the two fractions. (h) Line plot presents NTA data for the split sEV and non-sEV fractions. (i) Bar plot compares the signal produced by plasma, isolated sEV, non-sEV matrix, plasma (mimic) with a tenth of the non-sEV matrix, and plasma with ten times the non-sEV matrix. Error bars represent one standard deviation.

### Characterization of Small Extracellular Vesicles (sEVs) Using IJPs

To characterize small extracellular vesicles (sEVs), we employed a straightforward method that involves mixing IJPs with sample solutions and observing their blinking behavior under a fluorescent microscope, as illustrated in Fig. 2b. One significant advantage of using micron-sized IJPs is their ability to settle under gravity at a modest but noticeable rate. This settling allows the IJPs to move out of the microscope’s focal plane after measurements, continuously replaced by particles from the liquid above in the droplet. This process eliminates the need for microscopes equipped with z-stacking capabilities or microfluidic devices to supply fresh IJPs in-frame for measurements. Furthermore, this method facilitates extended observation periods, enhancing the statistical reliability of our measurements as more IJPs are analyzed. Typically, imaging across a 60-120 second duration proves sufficient for gathering statistically representative data. Additionally, given that the micron-sized IJPs generally exhibit a blinking period of 1-2 seconds, a standard camera with a frame rate of 10-30 frames per second can effectively capture this blinking. Therefore, the equipment required for IJP experiments is already available in most biological research laboratories. This ubiquity allows for the seamless integration of IJP methodologies into existing lab setups, enhancing their accessibility and facilitating adoption without requiring specialized equipment. It should be noted that the reported blinking periods represent the ensemble average across all IJPs.

Figure 2c shows the significant shift in blinking periods of IJPs with anti-CD63 capture when incubated with PBS (blank) compared to isolated 3E11 sEVs/mL derived from human plasma. This shift can be calibrated against known concentrations of isolated sEVs, as shown in Fig. 2d for different capture antibodies on IJPs, showing a limit of detection (LLOD) of 4 × 10^5 sEV mL⁻¹ (LoB = 5 × 10^4 sEV mL⁻¹) and > 3 log₁₀ dynamic range. The lower limit of quantification (LLOQ) is 1 × 10^6 sEV mL⁻¹, and the assay remains linear from 1 × 10^6 to 1 × 10^9 sEV mL⁻¹ on a semilog concentration–signal plot. Repeatability, intra-/inter-day variation (intra-day CV = 16%) and lot-to-lot precision (CV = 27%) are within the accepted range for sEV assays, and back-interpolated accuracy across three standard levels spans 87–101 %. In simulated plasma experiments, spike-in recovery is 104 % in both 10- and 1000-fold diluted, sEV-depleted matrices. Full validation data, including precision metrics, accuracy and recovery, are provided in Supplementary Table 1. Calculation was done using independently drawn calibration curve using anti-CD63+ IJPs. We have also shown robustness in the blinking time period distribution in Supplementary Fig. 1.

It is important to note that all antibodies for various proteins fall on the same universal curve for the IJPs when calibrated against the actual concentration of sEVs with that specific protein using Surface Plasmon Resonance. This occurs because we ensure a high probe concentration on the IJP surface with high-affinity antibodies, meaning that the reaction between sEV and capture antibody on the IJP is very fast (proportional to the product of surface coverage and the on-rate of antibodies); thus, they are limited by the diffusion of sEV from the bulk to the IJPs which provides the universal scaling and hence removes any antibody-to-antibody variation. Furthermore, this approach significantly reduces biases that might arise due to steric hindrance or avidity effects, particularly with sEVs that may have proteins adjacent to the target protein, exhibit slightly different conformations with exposed epitopes, or have a high density of surface proteins facilitating multivalent binding. These factors typically influence the kinetic on-rate and off-rate but are effectively neutralized under conditions where diffusion-limited mass transfer predominates, ensuring that the captured sEV fractions are truly representative of the bulk. This is discussed more in subsequent sections with further proofs.

We know that sEVs typically range from 50-200 nm in size, and therefore, using track-etched pores with no tortuosity allows us to filter out entities larger than the pore size from the biofluid. We applied this method to filter human plasma in three different ways and analyzed flow-through samples: The first strategy is with a 220 nm pore, which should not remove any sEVs; the second strategy is with a 100 nm pore, which should remove a significant number of sEVs larger than 100 nm but not all, and the third strategy is with a 50 nm pore, which removes all sEVs. As demonstrated in Fig. 2e, no significant signal is observed for the 50 nm filter, while the 100 nm filter shows a reduced signal compared to the 220 nm filter. This indicates that our platform is exclusively sensitive to larger structures, and entities smaller than 50 nm do not produce a signal. This is crucial as it selects against lipoproteins in plasma, a significant source of false signals in sEV assays, which are typically smaller than 50 nm. This unique size-dependent characteristic of our IJP assay platform enables the use of a simple pre-filtration step, in which the plasma samples are passed through a 220 nm porous membrane to remove any entities or interfering protein aggregates bigger than 220 nm, resulting in relatively cleaner samples for the assay.

Moreover, protein aggregates can potentially be a significant source of signal on the IJP surface. Fig. 2f demonstrates that using an isotype control under identical functionalization parameters produced no signal, effectively ruling out protein aggregates interacting with the IJP surface. Additionally, the treatment of sEVs with non-denaturing detergent, which does not lyse protein aggregates, resulted in no signal, as shown in Fig. 2f. This indicates that lysed sEVs release all their proteins into an almost soluble form due to the detergent disrupting the lipid bilayer. Yet these soluble protein counterparts, despite binding to IJPs, do not produce any significant change in blinking signal. This is essential information because, when IJPs are incubated in plasma, they encounter numerous proteins associated with sEVs. However, these same proteins could potentially be presented in a soluble form or on other lipoproteins. Therefore, the fact that entities smaller than 50 nm, which include both soluble proteins and lipoproteins, cannot produce a signal, thus confirming that IJPs can effectively address these challenges and potentially quantify sEVs directly from plasma.

To further demonstrate that the non-sEV component of plasma does not generate a signal, we utilized a recently commercialized NanoEx by Aopia Biosciences. This device efficiently separates plasma into sEV-containing and non-sEV matrices, as illustrated in Fig. 2g, with a non-tortuous asymmetric membrane (see Fig. 2h). This separation enables us to isolate plasma and create simulated plasma by adjusting the sEV to non-sEV ratio through recombination. In Fig. 2i, we show that simulated plasma with 10% or 1000% of the original non-sEV matrix/sEV ratio performs similarly to the plasma while maintaining the three samples maintained the same sEV concentration (see Methods for preparing simulated plasma). This finding indicates that the soluble fraction in plasma has a negligible effect on the signal, thereby making IJPs highly resistant to interference. It should be noted that signals may still be produced when an antibody cross-reacts to other proteins on sEVs, such as the L1CAM false-positive previously reported for sEVs^62^. Nevertheless, it falls within the scope of immunology to identify more specific antibodies, as non-specific antibodies can generate signals even in sandwich assays. The only known platform capable of handling such non-specific reactions is the proximity ligation assay, which faces its own challenges and requires extensive pretreatment due to using interference-prone Polymerase Chain Reaction (PCR) to generate signals. However, a significant advantage of our platform is that if an antibody non-specifically reacts with a protein not present on sEVs, it does not generate any signal. This is a notable improvement over platforms like Western Blot or Surface Plasmon Resonance (SPR), where signals are still produced when non-specific proteins are captured, whereas IJPs do not generate signals for entities smaller than 50 nm even if antibodies cross-react with non-sEV fraction.

### Characterizing the nature of the signal, blinking and normalization

In our study, we conducted a series of experiments using biotinylated nanoparticles of varying sizes to evaluate the performance of IJPs conjugated with anti-biotin antibodies to demonstrate the size dependence of blinking in a more controlled setting. Specifically, we utilized biotinylated beads of 20 nm, 50 nm, and 200 nm to represent different particle populations within our samples (Fig. 3a). The choice of anti-biotin over streptavidin for surface functionalization was deliberate as anti-biotin exhibits orders-of-magnitude lower affinity than the streptavidin– biotin pair^63^ Replacing it with streptavidin would introduce a tetrameric protein with exceptionally high avidity. This would complicate direct comparisons with the other monoclonal antibodies evaluated in this study, unlike anti-biotin, which also serves as a monoclonal antibody. Thus, the use of anti-biotin ensured consistency with our existing experimental protocols for antibody conjugation and highlighted the high sensitivity of our system. The results demonstrated that the IJPs exhibited a strong and specific binding to the biotinylated beads, with the signal intensity correlating with the size of the nanoparticles, with negligible signal with 20 nm nanoparticles, which would generally represent lipoproteins and smaller non-specific species in a biological setting. This finding underscores the robustness of our platform in detecting small extracellular vesicles (sEVs) and other nanoscale entities, while effectively excluding signals from smaller contaminants such as lipoproteins.

**Figure 3:**
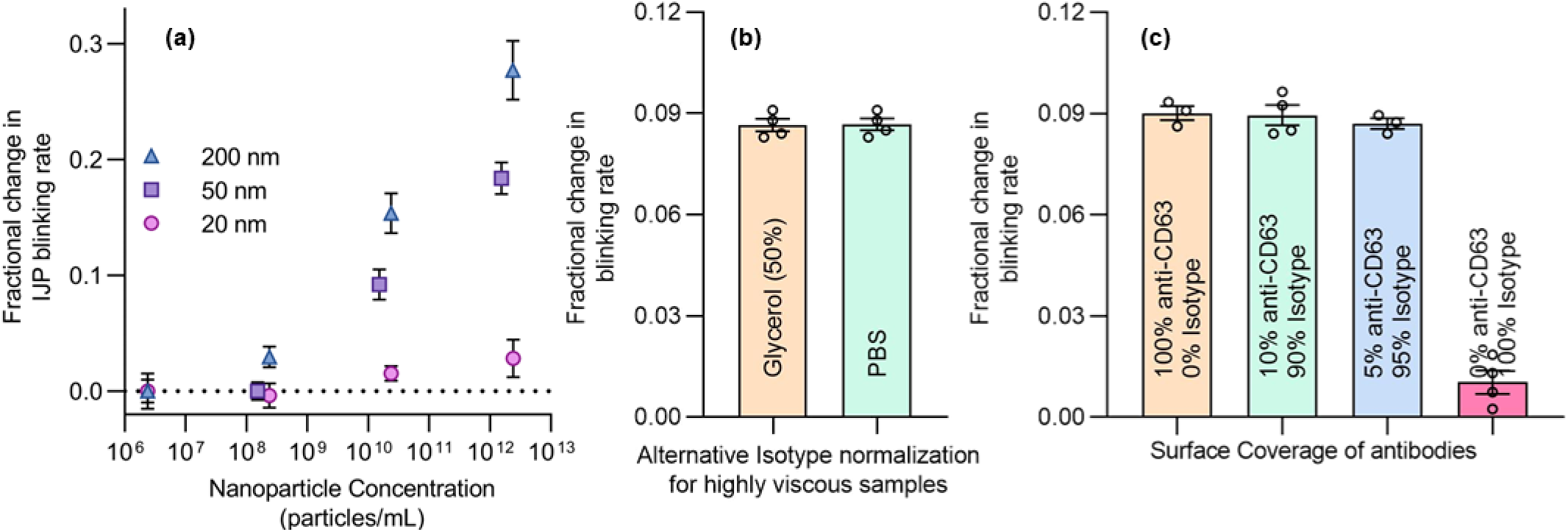
Illustrating the effect of particle size and surface marker coverage on the blinking rate of IJPs, highlighting the transport-limited nature of binding. (a) Shows the change in blinking rate with anti-biotin conjugated IJPs incubated with nanoparticles of different sizes. The calibration plot is slightly shifted from that for sEVs due to the highly negative zeta potential of these nanoparticles compared to the less negative sEVs trying to bind to the highly negative gold surface. (b) Alternative normalization for highly viscous samples: typically, normalization uses IJPs with an antibody to the surface marker incubated with a blank, but this approach uses isotype controls directly with the sample. (c) Competitive crosslinking alters antibody surface coverage, showing no correlation with surface coverage of target antibodies, suggesting transport-limited binding independent of on- rate, off-rate, or antibody concentration. This is evident from overlapping calibration plots with different antibodies using IJP against the true sEV concentration with that specific marker.

Furthermore, we investigated the effect of antibody density on the surface of IJPs revealed that the system operates in a diffusion-limited regime, where the observed response is largely independent of antibody affinity consistent with our universal calibration curve (Fig. 2d). This phenomenon is well-documented in our previous work^32^ as well as in techniques like Surface Plasmon Resonance (SPR)^64,65^, where mass transfer limitations can obscure the intrinsic kinetics of binding interactions. To validate this hypothesis, we employed a competition-based functionalization approach, introducing isotype antibodies as competitors to modulate the surface density of target-specific antibodies. Our results showed that varying the antibody density did not significantly affect the reaction rate, confirming that the system is dominated by diffusion rather than affinity. Only a negligible change in signal is observed, despite a 20-fold reduction in antibody concentration, thus confirming the binding is diffusion-controlled (Fig. 3c). This is very useful as it can allow for (a) a universal calibration plot with different antibodies targeting different proteins as long as the calibration is done against the true concentration of sEVs with that specific marker, and (b) same calibration plot for antibody targeting same protein but different epitopes of it. Additionally, no signal is produced with isotype, which confirms the high specificity of the signal.

Moreover, while our primary experiments demonstrated that 50x diluted plasma viscosity remains relatively stable and does not significantly impact the blinking rate of IJPs, we explored an alternative normalization approach to address potential viscosity-related variability in high-viscosity samples. By conducting experiments with IJPs functionalized with target antibodies and isotype control antibodies, we were able to normalize the blinking rates obtained from the isotype controls (as opposed to normal blank controls with IJPs with target antibodies), effectively removing the effects of nonspecific interactions and viscosity. This method proved effective even in samples resuspended in 50% glycerol, which has roughly ten times the viscosity of PBS (Fig. 3b). The consistency of our results across different viscosity conditions reinforces the robustness of our platform and its applicability to a wide range of biofluid samples.

### Direct Characterization of sEVs Across Various Biofluids Using IJPs, Orthogonal Comparisons, and Controls

This section outlines the characterization of different proteins on sEVs from various sources, including cell media (DiFi, GBM9, MDA-MB-468, A375P, and 3T3), human serum, and urine. The method produces protein concentrations comparable to those obtained through ultracentrifugation (UC) followed by Surface Plasmon Resonance (SPR) analysis, with both methods showing similar relative expressions. Measurements of CD63+, CD81+, CEA+, GPC1+, and aEGFR+ sEVs using IJP and UC+SPR, as shown in Fig. 4a and b, respectively, indicate identical trends and relative differential expressions. A challenge with SPR is the detection of soluble proteins even post-UC, hence double pelleting and resuspension were performed to ensure high purity of sEVs. The consistent results between the IJP platform, which takes under 60 minutes, and the UC+SPR platform, which requires almost a day, across various cell culture media as well as human serum and urine, suggest that IJPs can effectively and rapidly characterize sEVs with minimal interference.

**Figure 4:**
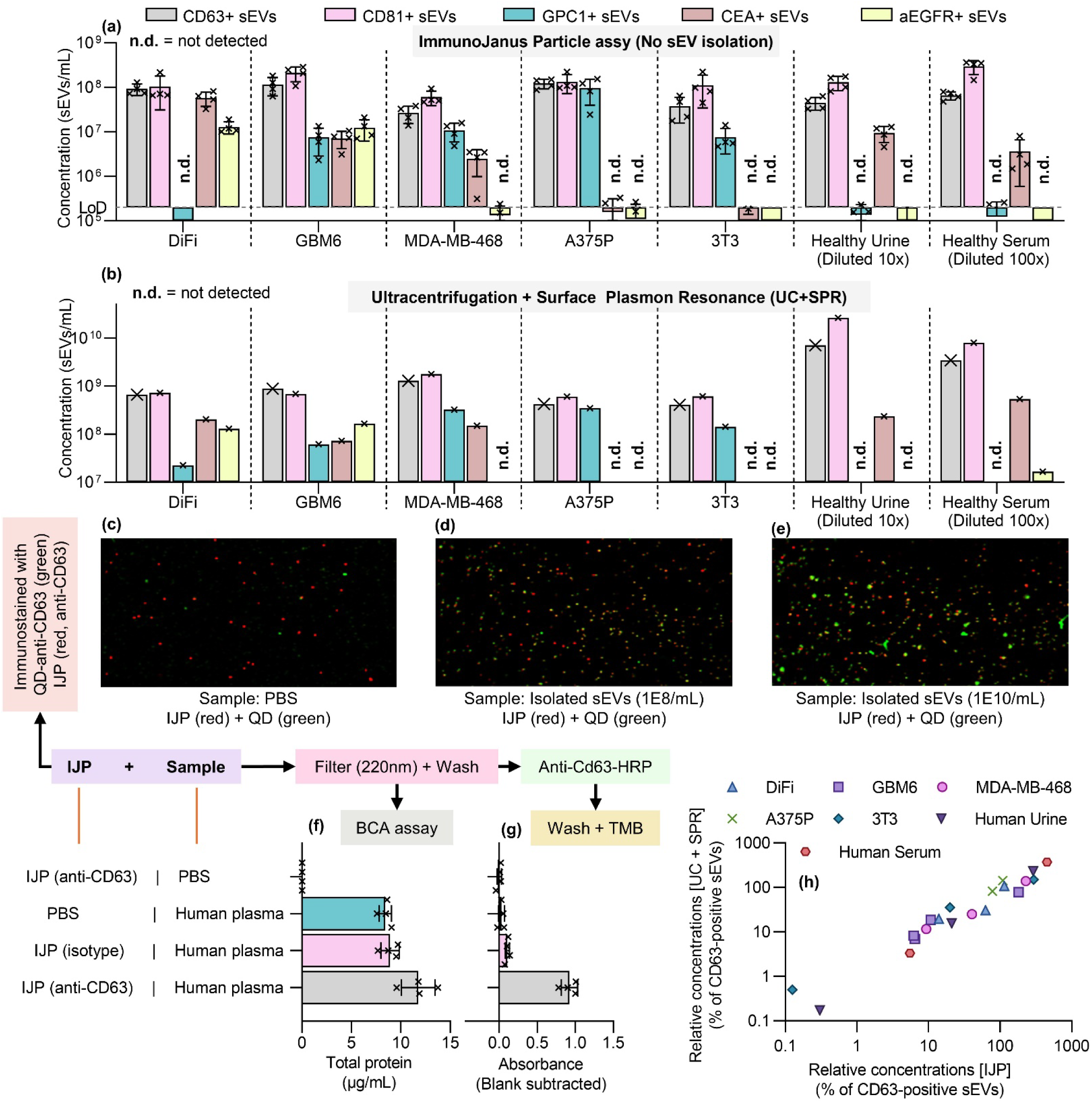
Comparing IJP characterization to orthogonal Ultracentrifugation (isolation) + SPR (characterization) (UC + SPR) and further controls for IJPs. (a) Characterization of CD63+, CD81+, GPC1+, CEA+, and aEGFR+ sEVs in cell media (DiFi, GBM6, MDA-MB-468, A375P, 3T3) and human biofluids (urine and serum) using IJPs. (b) Characterization of the same sEVs using UC + SPR, showing identical results. Panels (c- e) show immunostaining of IJPs (red) with anti-CD63 capture mixed with samples containing Quantum Dots labeled with anti-CD63 (green): (c) PBS, (d) 1E8 sEVs/mL, and (e) 1E10 sEVs/mL. (f) BCA assay of IJPs mixed with 10x diluted human plasma, filtered with 220nm, and washed with PBS, showing high non-specific binding of proteins for controls that do not change the blinking rate. (g) Instead of the BCA assay, anti-CD63 with HRP is added and washed, followed by an enzymatic reaction with TMB. IJPs producing significant shifts in blinking are highly enriched in anti-CD63, while those that do not change blinking significantly do not produce a high signal in this assay, despite having similar total protein concentrations as shown in (f). (h) UC + SPR and IJPs show a linear trend for the relative expression of different sEVs normalized by CD63+ sEVs. Error bars represent one standard deviation.

To verify that IJPs (red fluorescence) are capturing sEVs, we incubated anti-CD63 functionalized IJPs with PBS (control), 1E8 sEVs/mL, and 1E10 sEVs/mL, along with anti-CD63 conjugated quantum dots (QDs) (green fluorescence), as illustrated in Fig. 4c, d, and e, respectively. These figures demonstrate an increase in green fluorescence correlating with sEV concentration, underscoring the lower detection limits of fluorescence-based methods. Notably, QDs, known for their high fluorescence, produce weak signals at 1E8 sEVs/mL, whereas IJPs detect significantly lower concentrations of sEVs. Additional controls included incubating IJPs with samples, followed by mixing, filtering through a 220 nm filter, and performing a PBS wash, which were then analyzed for total protein and captured sEV. Specifically, total protein was measured using a bicinchoninic acid (BCA) assay (Fig. 4f) and anti-CD63 was detected using an enzymatic reaction with horseradish peroxidase (HRP)-conjugated anti-CD63 (Fig. 4g) on these captured beads. Fig. 4f indicates minimal non-specific binding to the IJPs (most on the 220 nm membrane based on control with no-IJP), while Fig. 4g shows substantial anti-CD63 binding in the positive controls, demonstrating the sensitivity and specificity of these assays. Due to the low sensitivity of the BCA and enzymatic immunoassays, a large volume of IJPs and samples was required to produce detectable signals, even with human plasma.

We analyzed the data from Fig. 4a and b, which measured sEV concentrations using the IJP platform and UC+SPR, respectively. We compared these results, normalized to CD63+ sEVs, and found that the IJP platform, without involving an isolation step, performed similarly to UC+SPR, as shown in Fig. 4h. This strong correlation indicates that IJPs can directly characterize a wide range of untreated biofluids, making the isolation step unnecessary when the characterization of sEV surface markers is the only objective.

### Rapid Screening of Disease States Using Small Extracellular Vesicles (sEVs) in human plasma

In this study, we introduce one of the initial applications of disease screening using small extracellular vesicle (sEV) surface markers, employing the IJP platform within a diverse cohort of human subjects. This pilot study includes individuals diagnosed with colorectal cancer (CRC), pancreatic ductal adenocarcinoma (PDAC), glioblastoma multiforme (GBM), Alzheimer’s disease (AD), and healthy controls. Our selection aims to reflect a realistic hospital setting where patients with various diseases coexist with healthy individuals, contrasting sharply with single-disease models that only incorporate one disease type alongside healthy subjects.

A primary challenge in global health is the accurate differentiation of healthy individuals from those with diseases and further pinpointing the specific disease. To address this, we selected four distinct sEV surface markers, each associated with specific diseases: conformationally active EGFR (mab806) for GBM, carcinoembryonic antigen (CEA) as a tumor marker for CRC and as a gastrointestinal tissue marker, glypican-1 (GPC1) for PDAC, and phosphorylated Tau181 (pTau181) for AD. These markers were chosen for their established links to these diseases, although the overexpression of markers in diseases other than those expected remains a significant screening challenge within a multi-disease cohort.

Our findings identified distinctive profiles of sEV populations: aEGFR+ sEVs were significantly elevated in GBM and CRC samples, but not in healthy, AD, or PDAC samples, as demonstrated in Fig. 5a. Conversely, GPC1+ sEVs were elevated in GBM, CRC, and PDAC (Fig. 5b), while CEA+ sEVs showed pronounced elevation primarily in CRC (Fig. 5c). Based on these observations, we developed a straightforward multi-disease screening protocol illustrated in Fig. 5d. Initially, we test for aEGFR+ sEVs, which stratifies our cohort into those with overexpression (GBM, CRC) and those without (Healthy, PDAC, AD), achieving an area under the curve (AUC) of 0.9899 and p-values around 2E-4 (Fig. 5e, i). For patients with overexpression (CRC or GBM), subsequent testing for CEA+ sEVs enables differentiation between GBM (no overexpression) and CRC (overexpressed CEA+ sEVs), with an AUC of 0.9602 and p-value ∼2E-4 (Fig. 5f, j). Conversely, if aEGFR+ sEVs are not overexpressed, we proceed with testing for GPC1+ sEVs, facilitating differentiation between PDAC (overexpressing) and Healthy/AD (no overexpression), with an AUC of 0.9279 and p-value ∼1E-2 (Fig. 5g, k). If GPC1+ sEVs show no overexpression, we then test for pTau181+ sEVs, which allows differentiation among healthy, AD, and mildly cognitively impaired (MCI) subjects, with an AUC of 1 between healthy and AD and a p-value of 5E-11 (Fig. 5h, l). This methodology offers a highly accurate approach for disease detection within a cohort using sEV markers characterized by IJPs.

**Figure 5:**
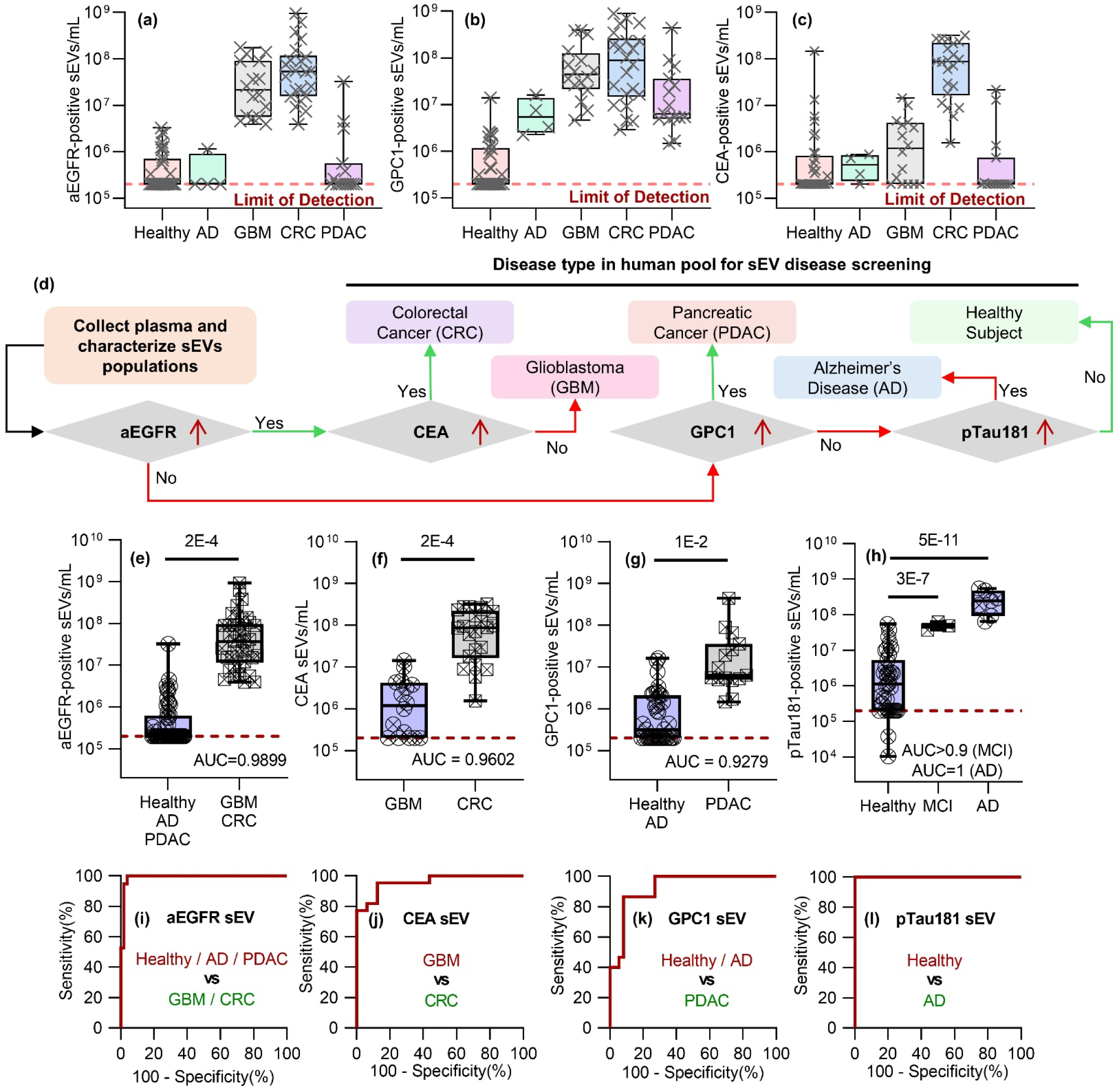
A blink disease screening 60-minute protocol using IJPs with 87 human subjects comprising healthy individuals, colorectal cancer patients (CRC), pancreatic adenocarcinoma patients (PDAC), glioblastoma patients (GBM), and Alzheimer’s patients (AD), using their plasma directly. (a-c) Typical sEV specific protein population profiling of human plasma for (a) aEGFR+ sEVs, (b) GPC1+ sEVs, and (c) CEA+ sEVs using box and whisker plot. (d) Schematic diagram illustrating the process of disease type screening in human plasma samples for small extracellular vesicles (sEVs). Plasma samples are collected, and sEV populations are characterized. The presence of specific sEV populations determines further testing with specific overexpressions of aEGFR+, CEA+, GPC1+, and pTau181+ sEVs yielding a corresponding disease. (e) Overexpression of aEGFR+ sEVs occurs only in GBM and CRC, not in Healthy, AD, and PDAC, ruling out multiple diseases. (f) Upon overexpression of aEGFR+ sEVs, testing for CEA+ sEV overexpression allows differentiation between subjects with GBM (underexpression) and CRC (overexpression). (g) If aEGFR+ sEVs are not overexpressed, GPC1+ sEVs are tested, with healthy and AD showing no overexpression but PDAC being highly overexpressed. (h) If GPC1+ sEVs are not overexpressed, pTau181+ sEVs are checked to differentiate between healthy individuals and AD patients. (i-l) ROC plots corresponding to the diagnostic performance of (e), (f), (g), and (h), respectively. (e)-(g) are represented using box and whisker plot with central line being the median, box being the 25th and 75th quantile, and whiskers representing 0th and 100th quantile.

To demonstrate the consistency of these results, we conducted orthogonal measurements of the patient samples using ultracentrifugation plus surface plasmon resonance (UC+SPR), with Figures 6a-c (heatmap showing mean expressioin in Fig. 6g) serving as orthogonal counterparts to Figures 5a-c, showing a similar trend across the disease groups. Due to yield bias between samples, we normalized all concentrations for UC+SPR against CD63+ sEVs, thus preventing direct comparison with IJP, which measures raw sEV concentration, whereas UC+SPR measures a normalized fraction. We also show that sEV associated cargo is better than the total cargo found in plasma with total CEA for healthy and CRC shown in Supplementary Fig. 2.

**Figure 6:**
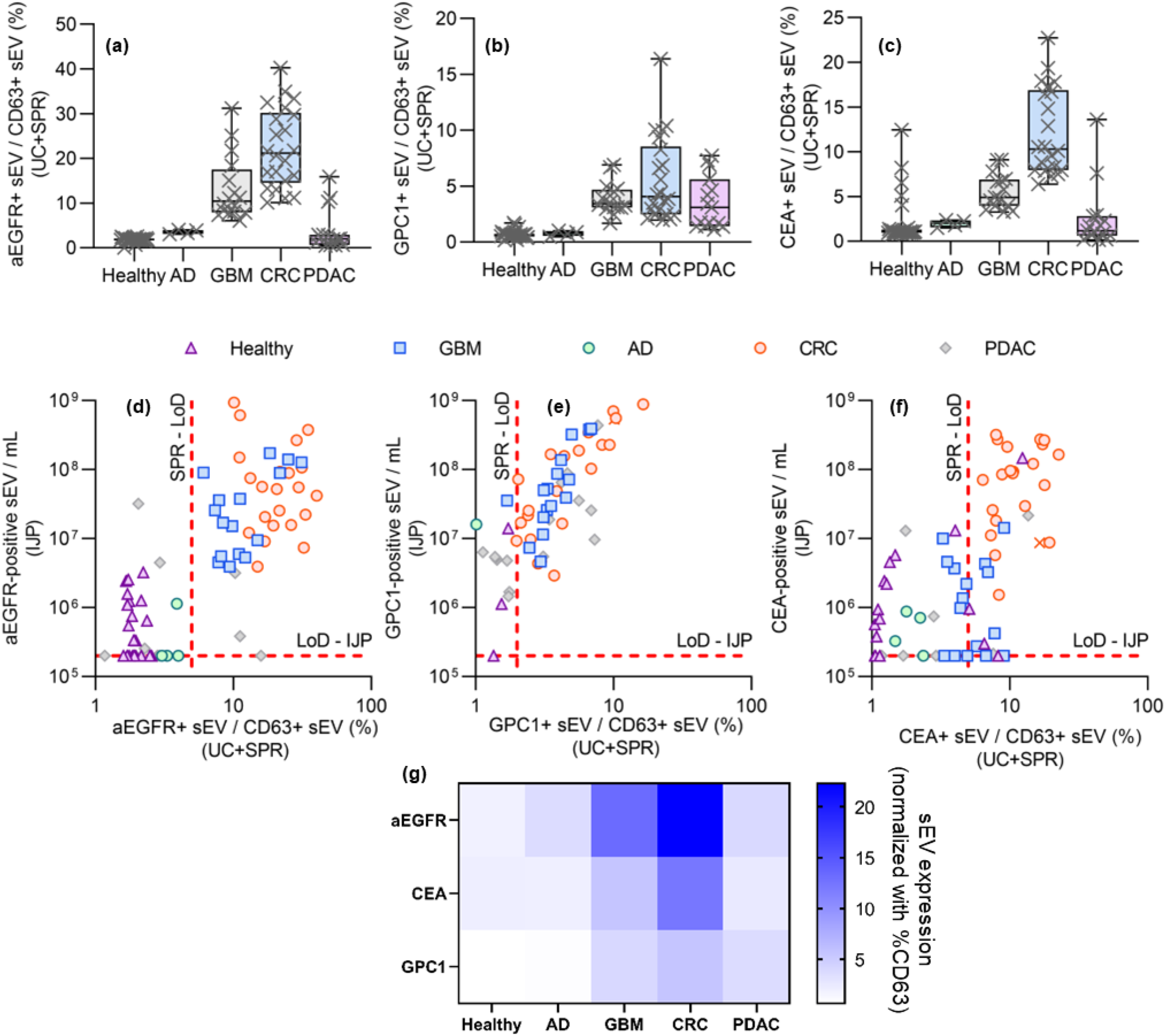
Orthogonal characterization for disease screening using sEV markers with UC + SPR. (a), (b), and (c) showing box and whisker plots correspond to the aEGFR+, GPC1+, and CEA+ sEV populations, respectively, using UC + SPR. The results and trends are consistent with those obtained using IJPs, with UC + SPR taking over 24 hours compared to less than one hour with IJPs. The box and whisker plot have a central line being the median, box being the 25th and 75th quantile, and whiskers representing 0th and 100th quantile. Panels (d), (e), and (f) show comparisons of the expression levels of aEGFR+, GPC1+, and CEA+ sEVs, respectively, across the human subjects when measured using IJP vs UC+SPR (normalized by CD63+ sEVs). The low sensitivity (several subjects below the limit of detection in SPR) and yield bias of ultracentrifugation, and the complexity associated with isolating relatively pure sEVs from human plasma compared to cell media, show a rough qualitative trend with clustering around the diagonal line with aEGFR+ sEVs showing moderate correlation (>0.3) and GPC1+ and CEA+ sEVs showing strong correlation (>0.5). Additionally, SPR is more prone to producing false signals from soluble proteins in non-vesicular fraction compared to IJPs that does not produce any signal for particles smaller than 50nm. (g) Heat map showing sEV expression (normalized by CD63) from different patient cohorts.

However, plotting the different proteins between IJP and UC+SPR for the three proteins (Figures 6d-f) also shows a similar trend, confirming that IJP is suitable for direct disease type detection with enhanced sensitivity and significantly reduced time requirement (under 60 minutes for IJP compared to almost a day for UC+SPR).

## Conclusions

In summary, this study introduces Immunojanus Particles (IJPs) as an effective tool for the direct characterization of small extracellular vesicles (sEVs) in biofluids, circumventing the need for prior isolation. Our results demonstrate that IJPs can accurately profile sEV surface markers with high sensitivity and specificity, directly from biofluid samples such as plasma, serum, and urine, and are robust against interference from dispersed proteins. The ability to rapidly detect and characterize sEVs from low volume samples not only offers a practical advantage over traditional methods but also improves the throughput and reliability of biomarker analysis in clinical settings. Unlike methods such as Nanosight, which analyze the overall population of sEVs and similarly sized nanocarriers, our platform provides high sensitivity for detecting sEVs expressing a specific surface marker. This targeted approach enables more precise and biologically relevant measurements.

The application of IJPs in a pilot study involving a mixed cohort of patients with different diseases and healthy controls underscores the method’s clinical relevance, particularly in the rapid screening of disease states. By providing a robust and streamlined approach to sEV analysis, IJPs hold potential for significant impacts in the fields of diagnostics and therapeutic monitoring, where quick and accurate biomarker assessment is critical. Further refinement and validation of this technology could lead to its adoption in routine clinical diagnostics, offering a non-invasive means for disease detection and management and a means for accelerating sEV research.

## Methods

### Ethical Statement

The studies involving human participants were reviewed and approved by the Indiana University Institutional Review Board (Study # 1105005445) and the Institutional Ethics Committee (IEC) at Austin Health, Melbourne, Australia. The patients/participants provided written informed consent to participate in this study. All ethical regulations relevant to human research participants were followed. We received Pancreatic Adenocarcinoma sample from Indiana Biobank, Colorectal cancer from Precision for Medicine, Alzheimer’s from Precision for Medicine and Indiana Biobank, Glioblastoma from Precision for Medicine from Andrew Scott and Hui Gan, Tumour Targeting Laboratory, ONJCRI, Melbourne, Australia and all healthy samples from Precision for medicine. All ethical regulations relevant to human research participants were followed.

### Immunojanus Particle Fabrication

The Immunojanus Particles were produced in-house using 1.0 μm FluoSpheres™ Polystyrene Microspheres from Thermo Fisher Scientific (Catalog no. F13081/F13083). The beads were diluted to 0.1% solids in 70% v/v isopropyl alcohol (VWR, Catalog no. BDH7999-4). A plain microscope slide (VWR, Catalog no. 48300-026) was treated with plasma for 15 seconds using an Electro-Technic Products Inc Model BD-20 High Frequency Generator. 1 mL of the dilute FluoSpheres™ suspension was deposited onto a slide and left for multiple hours – overnight to allow the isopropyl alcohol to evaporate completely. Once dried, the slide was inserted into an AIRCO Temescal FC 1800 electron beam vacuum deposition/thin-film coater system and coated with 30 nm of gold at a rate of 0.5 Å/s. The gold-coated slide was then removed and sonicated in an ultrasonic cleaner (Bransonic, Catalog no. 5510R-DTH) for 15 minutes. The released particles were collected in a 1% (v/v) Tween20(Sigma-aldrich P9416-50ML)/DI H2O solution and filtered three times using a 5 μm disk filters Cytiva Whataman™ Puradisc™, Catalog no. 10463533) to remove any aggregates and other large impurities. The particle suspension was concentrated to approximately 1 x 10^8 particles/mL and stored in a 4°C fridge until needed.

We used 1 x 10^7 IJP particles/mL for all experiments, as this particle concentration is the lowest concentration that can still provide quantitative capture and a reliable analytical signal. This rationale is based on our assumption that each 1μm IJP carries 10^4 antibodies, which corresponds to approximately 1 x 10^11 antibodies per mL or about 170 pM. The equilibrium dissociation constant of high-affinity monoclonal IgGs are in the 10-100 pM range^66^ The assay is thus operating above that range, ensuring that the binding is limited by vesicle availability rather than the antibody density. Therefore, 1 × 10^7 IJP particles per mL represents the lowest practical concentration that still provides quantitative capture.

### Antibody Functionalization on Immunojanus Particle

The gold hemisphere of the Janus Particles was functionalized with target specific antibodies using the Abcam Gold Conjugation Kit (ab154873). Up to 5 uL of antibodies underwent a buffer exchange using a 10K MWCO centrifugal filter (ThermoFisher Scientific, Catalog no. 88513) and 400 uL DI H2O. Antibodies were diluted to 0.1 mg/mL using the provided Abcam gold antibody diluent. 12 μL of dilute antibody was combined with 42 μL of the Abcam gold conjugation buffer in a 0.2 mL PCR tube (Axygen, Ref PCR-02-C). 45 μL of this solution was then combined with 50 μL of IJP solution in a PCR tube and mixed for 15 minutes at 1000 rpm on a shaker. Abcam gold conjugation quencher was added (5 μL) to the solution, which reacted for either 15 minutes at room temperature or overnight at 4°C. The functionalized IJPs were centrifuged in a Fisher AccuSpin Micro 17 at 6000g for 5 minutes and washed with 1:400 Tween20 (Sigma-aldrich P9416-50ML) DI H20 once, and twice with Dulbecco’s Phosphate Buffered Saline (Catalog no. 02-0119-1000) 10 times diluted in DI H20, before being reconstituted in 50 μL of 10x diluted PBS. This IJP solution was combined with samples in a 2:1 ratio and allowed to incubate for an hour before imaging.

In this research, the antibodies utilized included CD63 Mouse Monoclonal Antibody (Proteintech, Catalog: 67605-1-Ig, Lot: 10023876), CD81 Rabbit Polyclonal Antibody (Proteintech, Catalog: 27855-1-AP, Lot: 00107630), Glypican 1 Rabbit Polyclonal Antibody (Proteintech, Catalog: 16700-1-AP, Lot: 00056068), CEA Rabbit Polyclonal Antibody (Proteintech, Catalog: 10421-1-AP, Lot: 00017390), Mouse IgG1 Isotype Control Mouse Monoclonal Antibody (Proteintech, Catalog: 66360-1-Ig, Lot: 10028151) and Phospho-Tau181 Monoclonal Antibody (Invitrogen, Reference: MN1050, Lots: XF3582141, YI4023267). Mab806 antibody (ABT- 806, Catalog No. TAB-228CL) was purchased from Creative Biolabs, USA.

### Fluorescent imaging of Immunojanus particles

After incubating with the sample, a 2 μL drop of the IJP solution was pipetted onto a standard glass microscope slide (VWR, Catalog no. 48300-026). Three cover micro covers (VWR, Catalog no. 48366-089) are stacked on either side of the drop to create a vertical spacing of 440 μm. Another cover slip was placed on top of the solution so that contact with the drop was made. The entire setup was then placed on an Olympus IX-71 inverted fluorescent microscope above a 10x objective. An Olympus Optical Co, LTD 100 W High Pressure Mercury Burner (Model no. BH2-RFL-T3, no. 2308002) was used to create the fluorescence in the experiments. The focal plane was set to ∼220μm above the slide during recording. Videos ranging from 60s to 180s were captured using a Retiga EXi (QImaging, Catalog no. 01-RET-EXI-L-M-14-C) camera and a Basler ace 2 R (Basler, Catalog no. a2A1920-160ucPRO) camera at a frame rate of 10 Hz. All trials were conducted with a minimum of three technical replicates.

### Cell media preparation

DiFi cells, derived from human colorectal carcinoma, were cultured in a three-dimensional (3D) system to replicate the in vivo tumor microenvironment closely. The 3D scaffolds, constructed from type-I collagen at a concentration of 2 mg/mL, were layered in a tripartite structure: basal and top layers of pure collagen flanked a central layer embedding DiFi cells at a density of 5,000 cells/mL. This configuration was incubated at 37°C in a humidified atmosphere of 5% CO₂. Culture medium was supplemented with 10% fetal bovine serum (FBS), 2 μg/mL normocin, insulin-transferrin-selenium, epidermal growth factor, hydrocortisone, and T3 thyroid hormone, and refreshed every two to three days.

The human melanoma cell line A375P (RRID: CVCL_6233) and the human breast cancer cell line MDA-MB- 468 (RRID: CVCL_0419) were maintained in high glucose Dulbecco’s Modified Eagle Medium (DMEM, Gibco, USA) enriched with 10% v/v EquaFetal Serum (Atlas Biologicals, USA), 2 mM L-glutamine, 100 U/mL penicillin-streptomycin, and 1 mM sodium pyruvate. These cells were cultured under standard conditions at 37°C in a 5% CO₂ humidified atmosphere, ensuring optimal growth and maintenance.

Mouse fibroblast cells (3T3) were cultured in Minimum Essential Medium (MEM) supplemented with 10% FBS and 1% Antibiotic-Antimycotic Solution. The cells were housed at 37°C in a humidified 5% CO₂ environment. Passaging involved washing with 1X phosphate-buffered saline (PBS), trypsinization with trypsin-EDTA, and a recovery period of at least one day before experimental use. GBM9 glioblastoma cells were cultured as neurospheres in Neurobasal medium devoid of serum (Gibco) and supplemented with 3 mM GlutaMAX, 1x B- 27 supplement, 0.5x N-2 supplement, 20 ng/mL EGF (R&D Systems, MN), 20 ng/mL FGF (PEPROTECH, NJ), and 1% Antibiotic-Antimycotic Solution (Corning). Passaging was performed using the NeuroCult Chemical Dissociation Kit-Mouse (Stemcell Technologies, Canada) following the manufacturer’s guidelines.

### Immunojanus Particle immunostaining control

A quantum dot (Qdot) functionalization kit with an excitation/emission of 405/525 was first purchased from Thermo Fisher (Catalog no. S10449). The Qdots underwent conjugation according to the manufacturer’s protocol using a 100 ug cocktail of murine anti-human CD63 antibody (Proteintech catalog # 67605-1-Ig) and murine anti-human CD9 antibody (Proteintech catalog # 60232-1-Ig). IJPs functionalized with anti-CD63, as previously described, were mixed with purified sEVs of different concentrations for an hour and were then mixed with anti- CD63/anti-CD9 conjugated Qdot 525 for another hour. Then, the images were taken with Leica Stellaris 8 DIVE confocal microscope using a 10x objective. Two preset filters, the red FluoSpheres filter and the Qdot525 filter, were used in the sequential line scan mode to minimize spectral overlap, and maximize fluorescent yield. All the images are unprocessed, and full images for each channel are available in the supplementary data.

### Ultracentrifugation

The isolation and quantification of small extracellular vesicles (sEVs) from human plasma and concentrated cell media involve a detailed ultracentrifugation protocol. Initially, 200 µL of human plasma is diluted with PBS to a final volume of 1 mL, or 1 mL of concentrated cell media is used directly (concentrated using 100kDa filter from 10mL to 1mL). This mixture is centrifuged at 12,000 g for 20 minutes to remove larger particles. The supernatant is then passed through a 220 nm filter to eliminate larger debris. The filtered fluid is added on top of 3 mL of PBS in a 4 mL ultracentrifuge tube and ultracentrifuged at 167,000 x g for 1.5 hours using a swinging bucket rotor (Beckman Coulter SW60Ti), which is preferred for its efficiency in pelleting vesicles compared to fixed angle rotors. After ultracentrifugation, the supernatant is carefully removed, leaving about 0.2 mL to avoid disturbing the pellet, which is then resuspended in 0.5 mL of ice-cold PBS. For further purification, this resuspended solution is transferred on top of 15.5 mL of PBS in a 17 mL ultracentrifuge tube and ultracentrifuged at 167,000 x g for 4.5 hours (SW32.1Ti). This step aims to refine the sEVs by pelleting them again under high-speed centrifugation. After pelleting, the sEVs are resuspended and passed through a 300 kDa filter to remove soluble proteins, thus enhancing the purity of the sEV sample. This additional purification step ensures the isolation of high-quality sEVs, now ready for downstream analyses such as surface plasmon resonance (SPR) to study their composition or concentration.

### Surface Plasmon Resonance-based characterization of isolated sEVs

Prior to experimentation, all instrument-specific pre-experimental protocols recommended by the surface plasmon resonance (SPR) instrument manufacturer were followed, particularly those pertaining to "the maintenance chip" section. The system was then allowed to operate for an additional 12 hours using double-distilled water (DDI) in standby mode. Subsequent to this preparatory phase, the chip (Series S CM5 SPR chips, Cytiva, Catalog no. 29149603) was docked and normalized using 70% glycerol, or as otherwise advised by the manufacturer. The running buffer was then switched to phosphate-buffered saline (PBS) and maintained for two hours. Antibodies were buffer exchanged and resuspended in 10 mM MES buffer at a pH of 6.0. This solution was flowed over the SPR chip at a rate of 10 µL/min, delivering 100 µL of 0.1 mg/mL antibodies for 2 minutes to ensure appropriate preconcentration behavior. The selected flow channel was then prepared by flowing PBS until a stable baseline was achieved. To functionalize the channel, a coupling solution containing 20 mg each of EDC (1-ethyl-3-(3-dimethylaminopropyl) carbodiimide hydrochloride, Life Technologies, Catalog no. 22980) and Sulfo-NHS (N-hydroxysulfosuccinimide, Life Technologies, Catalog no. 24510) in 700 µL of 100 mM MES at pH 4.7 was flowed through the chip for 15 minutes at the same flow rate. Following this, approximately 400 µL of the antibody solution (0.1 mg/mL in 10 mM MES pH 6.0) was applied at 10 µL/min for 25 minutes. The reaction was quenched by flowing 0.1 M ethanolamine (Life Technologies, Catalog no. 022793.30) for 2 minutes. The channel was then washed with PBS until a stable baseline was reestablished. The extent of antibody functionalization was assessed by measuring the baseline shift before and after conjugation, with an expected increase of over 2000 response units (RU) indicating successful antibody coupling.

Baseline establishment and experimental conditions for surface plasmon resonance (SPR) measurements were conducted as follows: Initially, the baseline was recorded to ensure stability, defined as a drift rate of less than 0.1 response units (RU) per minute. If the drift exceeded this threshold, the running buffer was flowed overnight or until the baseline stability criteria were met.

Upon achieving a stable baseline, experimental procedures commenced. Two flow rates, 1 μL/min and 10 μL/min, were chosen to cover a decade of flow rate variation while minimizing sample volume consumption. At the start of each cycle, the system was first set to a flow rate of 10 μL/min: the running buffer was flowed for five minutes, followed by a sample injection in high-performance mode for two minutes. Subsequently, the system was washed with the running buffer for 60 seconds before sequentially injecting Glycine-HCl (10 mM, pH 2) and 1% albumin, each for 20 seconds. The flow rate was then reduced to 1 μL/min, and the running buffer was flowed for an additional 30 minutes, followed by a five-minute sample flow. The flow rate was subsequently restored to 10 μL/min for final wash steps and injection sequences identical to the initial set. For both flow rates, the slope of the sensor response between 30-120 seconds post-injection was recorded to assess binding characteristics.

To correct for any system artifacts, a blank cycle was performed immediately following the sample measurements, using phosphate-buffered saline (PBS) instead of the sample to simulate identical injection conditions. The process was identical to that of the sample injections, including the adjustment of flow rates and the sequence of buffer and reagent injections. The measured slopes from the sample and blank cycles were compared to ascertain the specific binding response. These steps were repeated for each new sample, with the resultant signal representing the differential between the slopes obtained during sample and blank measurements, thus providing a corrected and reliable measure of the binding interactions. Based on the mass transfer constant in a laminar flow, size of sEVs, size of sensor chip and diffusivity of sEVs, they are converted to sEV number concentrations.

### Creating simulated plasma with varying non-sEV proportions at equivalent sEV concentrations

To produce simulated plasma, we first isolated the small extracellular vesicles (sEV) and non-sEV matrix from 50-fold diluted plasma. This process yielded quantities of sEV and non-sEV matrix equivalent to those found in the original volume of diluted plasma. Specifically, we used 𝑉𝑉_1_ mL of 50-fold diluted human plasma to generate

𝑉𝑉_2_mL of sEV and 𝑉𝑉_3_mL of the non-sEV matrix. To maintain a constant sEV concentration, we aimed to reproduce a specified percentage 𝑥𝑥(%) of the original non-sEV fraction while keeping the sEV fraction unchanged. We fix sEV concentration to the amount present in 1000-fold diluted plasma, including for the 50- fold plasma, by diluting it 20 times.

For creating 𝑉𝑉_0_ mL of a simulated sample that mimics the characteristics of 1000-fold diluted plasma, we mixed the following components in specified proportions:

For 10% of the original non-sEV matrix fraction in proportion to sEV (1:0.1 sEV/non-sEV matrix in Fig. 2i)

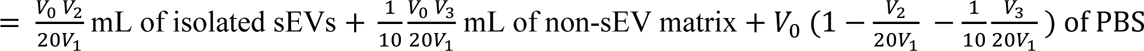

For 1000% of the original non-sEV matrix fraction in proportion to sEV (1:10 sEV/non-sEV matrix in Fig. 2i)

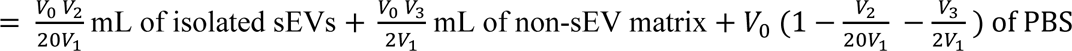

## Acknowledgement

This work was partially supported by the NIH Commons Fund, through the Office of Strategic Coordination/Office of NIH Director, 1UH3CA241684-01 (HCC & SS). We are also grateful for cell media samples from Robert Coffey, Al Charest, Yichun Wang and Crislyn D’Souza-Schorey. We would like to acknowledge Notre Dame Integrated Imaging Facility (NDIIF) at Notre Dame. We would also like to *thank the Notre Dame Biophysics Instrumentation Core Facility* for use of the Biacore T200 SPR system, purchased with funding from NIH (S10 OD028553). SK would like to acknowledge O’Brien Fellowship from Berthiaume Institute of Precision Healthy (BIPH) at Notre Dame.

## Author contributions

HCC and HSC conceived the project. SK and JAS designed the experiments, ran the IJP experiments, analyzed experimental results, and completed data processing. SK and JAS also did the UC+SPR and made all the figures for this paper. SS and JAS optimized functionalization of IJPs, S.K and JAS optimized the entire IJP workflow. TS, SK and JAS performed NTA. SK and HCC optimized the code for automated IJP blinking detection. All authors contributed to writing the manuscript.

## Data Availability

All data supporting this study in the main manuscript or the supplementary data.

## Code Availability

All primary codes critical to this study provided in the supplementary data.

