## Supplementary Information for "Immunojanus Particles for low-volume and isolation-free unlabeled characterization of small Extracellular Vesicle in biofluids: Characterization of disease type by surface marker profiling"

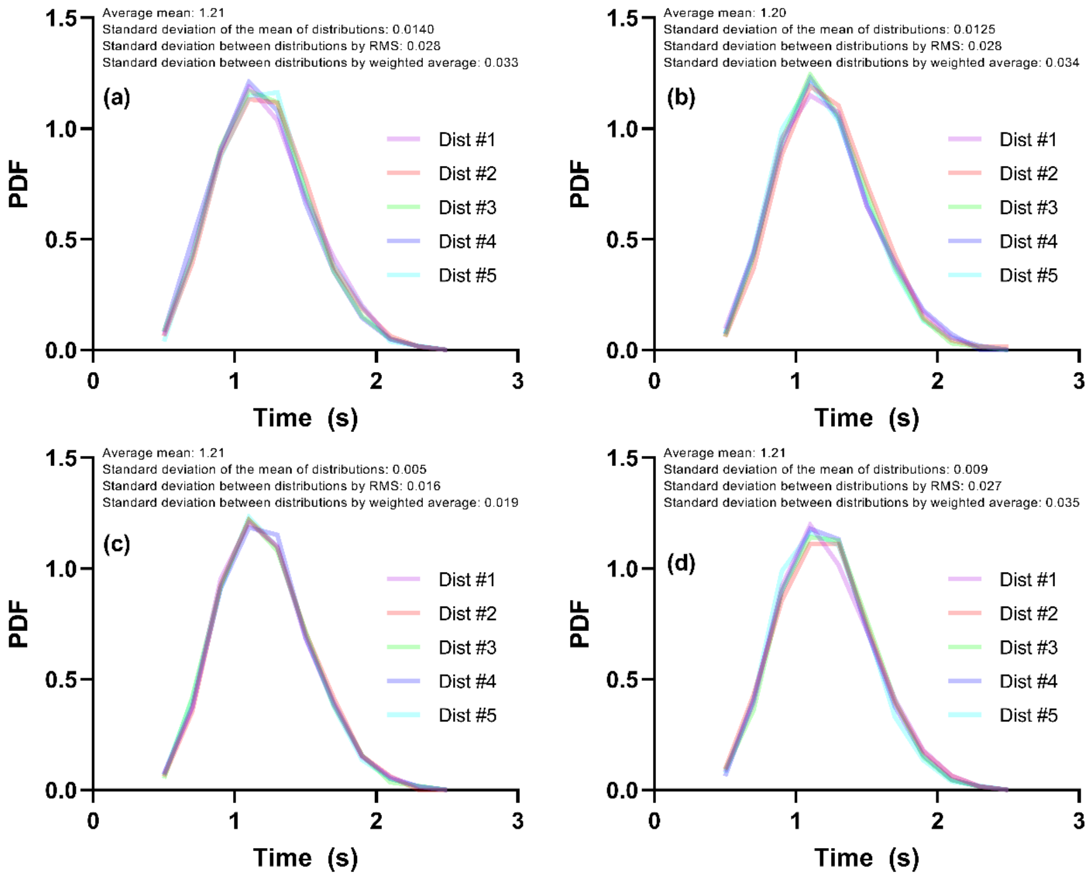


Supplementary Figure 1: Blinking time period distribution of IJPs from four different batches of IJP as shown in (a)-(d). Each contains approximately 300 recorded blinking IJPs in the focal plane of the microscope during recording and measurement. Each batch shows five different distributions that were functionalized with antibodies independently.

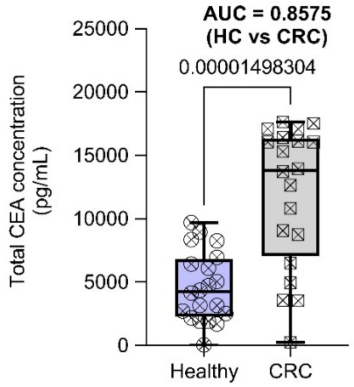


Supplementary Figure 2: Total CEA in human plasma across healthy and CRC cohorts

*Supplementary Table 1: IJP platform assay parameters. sEV stability during storage should be considered coupled with these numbers, and these numbers can be significantly reduced by averaging over more blinking IJPs.*

| **Metric** | **Method** | **Result** |
|---|---|---|
| LoB | Concentration at ~ Mean of blank + 1.65 x $\sigma$ (blank) | 5E4 sEV/mL |
| LLOD | LoB + 1.65 × $\sigma$ (blank) | 4E5 sEV/mL |
| LLOQ | Lowest point in calibration curve where CV<20% | 1E6 sEV/mL |
| Intra-day precision | Same sEV aliquot measured on same IJP batches | CV = 16 % |
| Lot-to-lot precision | Different sEV aliquot (same biologic replicated) measured on different IJP batches | CV = 27 % |
| Back-interpolated Accuracy | Measuring known standards and measuring accuracy as % of original concentration measured. | 97% (3E8 sEVs/mL)<br>87% (3E7 sEVs/mL)<br>101% (2E6 sEVs/mL) |
| EV spike recovery | Final concentration of 5E7 sEVs/mL spike<br>in 10x diluted plasma matrix (sEV depleted)<br>in 1000x diluted plasma matrix (sEV depleted) | <br>104%<br>104% |

Note – all these parameters were calculated for > 200 blinking IJPs per replicate. These numbers can be different if averaged over fewer IJPs.
